# Combination of alpha-synuclein aggregation inhibitor anle138b and ER stress inhibitor AMG PERK 44 increases neuroprotection in Parkinson’s disease organoid model

**DOI:** 10.64898/2026.03.16.712219

**Authors:** Natalia Siwecka, Michał Golberg, Grzegorz Galita, Ireneusz Majsterek

**Author notes:** Corresponding author. E-mail address (I. Majsterek).

## Abstract

Parkinson’s disease (PD) is the second most common neurodegenerative disease, resulting from accumulation of α-synuclein (α-syn) in midbrain dopaminergic neurons and progressive neuronal loss. The most relevant species of α-syn, oligomers, may exert neurotoxicity in a variety of mechanisms. Accumulation of misfolded α-syn in the endoplasmic reticulum (ER) lumen induces ER stress conditions that leads to activation of the Unfolded Protein Response (UPR) and its main sensor PKR-like ER kinase (PERK). PERK is critical for cell fate determination – under prolonged ER stress, it may direct cell towards pro-apoptotic pathways. Targeting of α-syn aggregation or UPR by genetic and pharmacological approaches proved effective in preclinical models of PD by previous research. Thus, in the present study, we aimed to determine the potential effect of combination of small-molecule inhibitors of α-syn aggregation and ER stress-mediated PERK signaling (namely anle138b and AMG44) in a novel, 3D in vitro model of PD. We demonstrate that combination of both anti-aggregation and ER stress-targeting approaches amplifies neuroprotection against PD in organoid model in terms of increased neuronal metabolic activity, decreased α-syn phosphorylation and aggregation, reduced dopaminergic cell death, and restoration of proteostasis.

## 1. Introduction

Parkinson’s disease (PD) is the second most common neurodegenerative disorder after Alzheimer’s disease (AD) and the most frequent synucleinopathy. PD affects over 10 million individuals worldwide and about 1% of the population over the age of 60 [1, 2]. As PD remains incurable with the symptoms intensifying with the disease progression, it is of utmost importance to improve PD patients’ quality of life. Currently, the only available treatment options for PD are symptomatic and, to date, there is no available neuroprotective or neurorestorative therapy for PD [3]. The pathophysiology of PD is complex and not fully understood, and thorough understanding of what really causes PD is a key to development of new optimized therapies [4].

The major molecular event underlying the pathophysiology of PD is the abnormal accumulation of α-synuclein (α-syn) within the dopaminergic neurons of the midbrain, that contributes to progressive neuronal loss [5]. Numerous factors, including mutations or multiplications of *SNCA* gene encoding for α-syn, and several post-translational modifications may disrupt the balance between α-syn production and clearance and make it prone to aggregate [6, 7]. α-Syn may misfold into various forms, from oligomers, through fibrils, to eventually assemble into Lewy bodies (LBs) [8]. Among all forms of α-syn, the oligomeric species are considered the most relevant as they may exert neurotoxicity in a variety of mechanisms, impair synaptic function, self-replicate and propagate from cell to cell in prion-like fashion [9]. Multiple lines of evidence have suggested that ER stress triggered by oligomers is particularly implicated in the pathogenesis of PD. Accumulation of misfolded α-syn within the ER lumen induces ER stress conditions that leads to activation of the Unfolded Protein Response (UPR) [10–13]. The UPR consists of three major stress sensors, inositol-requiring enzyme 1 (IRE1), activating transcription factor 6 (ATF6) and PKR-like ER kinase (PERK). The UPR branches are activated upon dissociation of binding immunoglobulin protein (BiP) once it detects the misfolded protein [14]. Consistently, it was found that α-syn may activate UPR directly via interaction with BiP chaperones [15, 16]. Among the three UPR sensors, PERK is critical for determining cell fate in PD [17, 18]. Activated PERK phosphorylates eukaryotic translation initiation factor 2 subunit 1 (eIF2α), that aims to restore proteostasis via transient inhibition of protein synthesis and selective upregulation of target pro-survival genes. If this approach fails, the cell is targeted to programmed cell death via activation of PERK-dependent, pro-apoptotic C/EBP-homologous protein (CHOP) [14]. Prolonged ER stress leads to mitochondrial damage and cell death in a positive feedback loop, via generation of ROS and calcium leakage that further potentiate α-syn accumulation and cellular impairment [14, 19, 20]. There is increasing evidence that UPR may be the main pathway involved in α-syn-related toxicity, as targeting of UPR by genetic or pharmacological approaches proved effectiveness in preclinical models of PD [11, 21–23]. What is more, increased levels of UPR markers, p-PERK and p-eIF2α, were detected in brain specimens from PD patients as compared to controls [18, 21].

Recent disease-modifying attempts for PD focus on different aspects of α-syn aggregation process, the most optimal approach of which is targeting of toxic oligomers [24]. Recently tested protein-based therapeutics like vaccines, antibodies or chaperones might have difficulty in passing through blood-brain barrier (BBB) and trigger collateral immunological reactions, hence small molecules targeting α-syn seem to be better candidates for treatment of brain diseases [25]. One of the most promising drugs is anle138b, a diphenyl-pyrazole (DPP) derivative discovered via high-throughput screening combined with medicinal chemistry optimization. Among a number of 10,000 compounds tested, anle138b appeared to be the most effective, of excellent oral bioavailability and BBB penetrance, and innocuous to treated animals [26]. Anle138b is an oligomer modulator that specifically inhibits the formation and accumulation of α-syn oligomers and acts on a broad spectrum of other amyloidogenic proteins that often co-aggregate. The compound was tested in multiple rodent models of prion disease, AD, PD, and multiple system atrophy (MSA) with astonishing results [26–28]. Not only did anle138b provide significant neuroprotection, but it also slowed the disease progression, even when the treatment started after the onset of symptoms or at the late-stage, which is important because most cases of PD are diagnosed at these stages [26]. The molecule is currently completing phase I clinical trials. Although the compound showed excellent results in several animal models, little is known regarding its effectiveness in human-based cellular environment.

Among UPR-targeting inhibitory molecules, the compound 44 developed by Amgen (AMG44) is a highly potent PERK inhibitor with an IC50 of 6 nM, demonstrating significant selectivity over other kinases like GCN2 and B-Raf. AMG44 is also characterized by oral bioavailability and good pharmacokinetic profile [29]. In contrast to other known PERK inhibitors, AMG44 does not interact with the receptor-interacting serine/threonine-protein kinase 1 (RIPK1), thus it is regarded as a useful tool for selective pharmacological PERK inhibition [30]. As AMG44 does not induce pancreatic toxicity [31], it could be a promising alternative to previously developed series of GSK inhibitors of PERK which, although effective in PD in vivo studies, exerted severe adverse events in tested animals [21]. Importantly, there is growing evidence that either long-term inhibition of p-eIF2α or its chronic activation can lead to substantial cytotoxicity, and therefore partial inhibition or activation of eIF2α phosphorylation would be an optimal strategy [32]. We have previously evaluated the neuroprotective properties of AMG44 in neurotoxin-based in vitro model of PD, in which it provided neuroprotection inter alia by fine-tuning of eIF2α activity [33].

What is more, it is suspected that too intensive disruption of protein aggregates or inhibition of oligomer transition to more stable, insoluble deposits could lead to ER stress and cell damage [34, 35], and we hypothesized that this effect should be effectively compromised by addition of ER stress inhibitor [36]. It has also been reported that anle138b failed to fully restore neural function in acute, neurotoxin-based MSA model, which provides rationale to combine it with the other compound, somewhat more specific to oxidative stress-related apoptosis [37]. Combination therapy possesses significant advances over single-agent therapy for it increases treatment efficacy, prevents development of drug resistance, reduces drug dosage, duration of treatment and decreases risk of adverse effects [36]. Inhibition of both accumulation and propagation of α-syn oligomers and the pro-apoptotic, UPR-dependent signaling pathways by the small-molecule inhibitory compounds may emerge as a first innovative, disease-modifying strategy, and the results could significantly contribute to research in PD field.

Considering the above, the main aim of the present research was to determine the potential effect of the small-molecule inhibitors of protein aggregation and ER stress-mediated PERK signaling against neurodegeneration in a novel, 3D in vitro model of PD. A recent development of organoid models is a major technological advance in preclinical studies, as they constitute a bridge between traditional 2D in vitro models and in vivo models [38]. Here, we developed a new model of sporadic PD – human midbrain organoids generated from induced pluripotent stem cells (iPSCs) treated with α-syn pre-formed fibrils (PFFs) and neurotoxin 6-hydroxydopamine (6-OHDA). This model is able to reflect human disease more accurately than widely used 2D cultures or animal models, and it demonstrates key pathological hallmarks of PD - α-syn aggregation and neuronal cell death. Animal models widely used in PD research possess several serious limitations – most of them bypass molecular events associated with natural course of human PD, and thus they cannot be taken as a translational proof-of-concept [39]. Moreover, use of cell-based in vitro models aims to relieve animals from unnecessary harm and distress. The selected small-molecule inhibitory compounds anle138b and AMG44 were investigated in PD organoids in terms of cytotoxicity, effect on α-syn accumulation, the level of apoptosis, and the modulation of the expression level of ER stress markers.

## 2. Methods

### 2.1 Generation of a 3D in vitro model of PD

As midbrain (mesencephalon) is the main brain part affected in the course of PD, in this study we utilized human midbrain organoids to model PD features in vitro. The organoids were generated from the normal human iPSC line, specifically Cellartis® Human iPS Cell Line 18 (ChiPSC18) purchased from Takara Bio (Cat. No. Y00305). These cells were obtained by reprogramming skin fibroblasts from a healthy male donor and are capable of differentiation towards neuronal fate. Indicated iPSC line was initially cultured in the dedicated Cellartis DEF-CS Culture System according with the vendor’s instructions, then the culture conditions were switched to mTeSR™1 medium (STEMCELL Technologies) for the next ≥2 passages. iPSC colonies were maintained daily with the differentiated areas regularly removed and passaged at 70% confluency (day ∼7) in order to ensure an optimal quality and pluripotency. High-quality iPSC colonies (below 10 passage) underwent differentiation according to the protocol provided by Paşca et al. [40]. Briefly, the iPSC were dissociated and seeded into microplate wells to form the embryoid bodies (EBs), and then subsequently cultured in the organoid formation medium, expansion medium and differentiation medium until day 43, when the organoids were transferred to final maintenance medium in the ultra-low attachment 6-well plate for culture maintenance of mature organoids. To confirm the midbrain-specific differentiation, the organoids were analyzed for the presence of specific cell markers: neuronal progenitors (FOXA2), neurons (MAP2, NFP) dopaminergic neurons (TH), and astrocytes (S100β) at day 30 and 55 by the immunofluorescence as described below. In addition, the level of specific neuronal markers (TH, Tuj1) and α-synuclein (SNCA) in the midbrain organoids was also evaluated by Western blot as described below. The midbrain tissue morphology within the organoids was additionally confirmed by histological hematoxylin/eosin staining.

For the induction of α-syn pathology, the normal human midbrain organoids were seeded with α-syn PFFs. PFFs are similar in structure to the building blocks of LBs, they trigger accumulation of endogenous α-syn and initiate PD-specific molecular perturbations; this model exhibits course of the disease close to that of human condition. The human α-syn PFFs and monomers were obtained from StressMarq Biosciences. Before application, α-syn PFFs were sonicated (30 cycles of 30s on/off on high power setting) and controlled for amyloid formation by ThT binding assay. ThT is a fluorescent dye that binds to beta sheet-rich structures, like those in α-syn PFFs. To perform the assay, 10 μM α-syn monomer, 1 μM α-syn PFFs or both were incubated with 25 μM thioflavin T (Sigma) in a total reaction volume of 100 μl at 37°C with shaking at 600 rpm for 10 min in the dark; than the fluorescence was measured with excitation/emission of 450/485 nm using a Synergy HT microplate reader.

Additionally, to accelerate neurodegeneration, the organoids were treated with 6-OHDA (Sigma-Aldrich), which significantly affects cell viability, induces oxidative stress, cell death and amplifies α-syn aggregation. While PD organoids were treated with α-syn PFFs (4 µg/ml) and 6-OHDA (300 µM) for 48 h, the control organoids were incubated with the equivalent amount of 0.1% DMSO and α-syn monomers for 48 h prior to detection. The use of both α-syn PFFs and 6-OHDA mimics the two key aspects of PD pathogenesis, namely α-syn aggregation and dopaminergic cell death. A 48 h incubation period was proven sufficient for induction of α-syn aggregation and partial cell death without causing extensive loss of viability in organoids. All applied chemicals are listed below in Table 1.

**Table 1.**
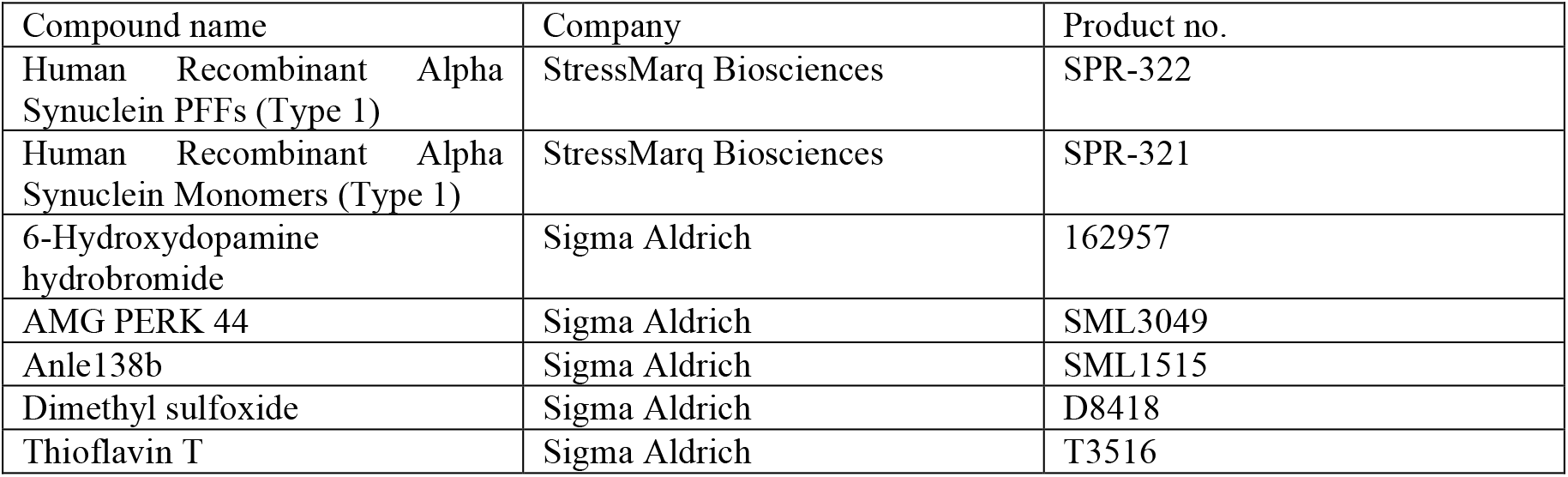
The full names and catalog numbers of the reagents applied in the experiments.

### 2.2 Assessment of cell viability

The cytotoxicity analysis of small-molecules anle138b and AMG44 in midbrain organoids was performed by CellTiter-Glo 3D viability assay (Promega). This assay is based on ATP quantification, which indicates the presence of metabolically active cells within the organoid. Each experiment was performed on the organoids in the same batch and of similar diameter. The organoids were transferred to white 96-well plates (Greiner) pre-treated with Anti-Adherence Rinsing Solution (STEMCELL Technologies) and incubated with each compound separately in a range of concentrations (0.1-200 µM) for 24 h. Then, the CellTiter-Glo 3D reagent was added, the samples were shaken at 900 rpm for 5 min and incubated for 25 min at room temperature (RT), away from light. Following the incubation, the luminescence was recorded and normalized to the mean luminescence of 0.1% DMSO-treated controls within the same plate. The luminescence was detected using a Synergy HT microplate reader (BioTek).

After confirmation that none of the applied inhibitors is cytotoxic towards normal organoids, the effect of each drug alone on the viability of ∼60-day PD organoids was tested at concentrations of 0.1-200 µM and at 24 h post-treatment using the CellTiter-Glo 3D assay. Then, the combination of drugs at their most effective concentrations (post-treatment with AMG44 at 25 µM and anle138b at 25 µM) was evaluated in PD organoids by the same method.

### 2.3 Immunofluorescent detection of α-syn

To determine the inhibitory effect of the compounds on the formation of α-syn deposits (aggregated α-syn, α-syn phosphorylated at Ser129 and total α-syn) the immunofluorescence staining was applied. PD organoids at day ∼60 were treated with the compounds in one of the following patterns: cell culture medium supplemented with AMG44, anle138b, or with both AMG44 and anle138b. Following the treatment, the organoids were fixed in 10% neutral buffered formalin solution supplemented with protease and phosphatase inhibitor cocktail (Roche) overnight, and embedded in paraffin, forming a formalin-fixed paraffin-embedded (FFPE) block. Microsectioned at 4 µm thickness samples were transferd to glass slides. Slides were than deparaffinized, rehydrated, and underwent antigen retrieval process. Then, the slides were blocked (3% BSA in PBS for 60 min) for 1h at RT and incubated overnight at 4°C with primary antibodies against pSer129 (Cell Signaling Technologies), agSyn (Abcam) and total-SYN (Invitrogen). On the next day, the sections were incubated with AlexaFluor secondary antibodies (Cell Signaling Technologies, Life Technologies) for 1 h at RT and ounterstained with Hoechst (1:2000) for 10 min away from light. Then, the samples were mounted and images were captured by fluorescent microscope (Nikon). Quantification of each staining was normalized to nuclei (Hoechst). All used antibodies are listed in Table 2 below.

**Table 2.** The full names and catalog numbers of the applied antibodies and fluorescent dyes.

| Target | Antibody name | Dilution | Company | Catalog No. |
| --- | --- | --- | --- | --- |
| FOXA2 | FoxA2/HNF3β (D56D6) XP® Rabbit mAb | 1:400 | Cell Signaling Technology | 8186 |
| MAP2 | Monoclonal Anti-MAP2 antibody produced in mouse | 1:400 | Sigma-Aldrich | M4403 |
| NFP | Neurofilament Protein, Clone 2F11, FLEX RTU. Monoclonal Mouse Anti-Human, Ready-to-use antibody | RTU | Dako | GA60761-2 |
| TH | Anti-Tyrosine Hydroxylase Antibody | 1:400 | Sigma-Aldrich | AB152 |
| S100 | S100, FLEX RTU. Polyclonal Rabbit Anti-, Ready-to-use antibody | RTU | Dako | GA50461-2 |
| Aggregated α-synuclein | Rabbit monoclonal [MJFR-14-6-4-2] to Alpha-synuclein aggregate - Conformation-Specific | 1:1000 | Abcam | ab209538 |
| pS129 α-synuclein | Phospho-α-Synuclein (Ser129) (D1R1R) Rabbit mAb | 1:500 | Cell Signaling Technology | 23706 |
| α-synuclein | alpha Synuclein Monoclonal Antibody (Syn 211) | 1:500 | Invitrogen | 32-8100 |
| Rabbit Ab | Donkey Anti-Rabbit IgG H&L (Alexa Fluor® 647) | 1:500 | Abcam | ab150075 |
| Mouse Ab | Goat anti-Mouse IgG (H+L) Cross-Adsorbed Secondary Antibody, Alexa Fluor™ 488 | 1:1000 | Invitrogen | A-11001 |
| nuclei | Hoechst 33342 solution, 20 mM | 1:2000 | Thermo Scientific | 62249 |

### 2.4 Cytometric apoptosis analysis

Analysis of the level of apoptosis of dopaminergic neurons in PD organoids treated with the compounds was evaluated by flow cytometry (FC) using FITC-conjugated Annexin V (AV)/propidium iodide (PI) staining (BD Pharmingen). This method is based on the affinity of AV to bind phosphatidylserine at the outer membrane of apoptotic cells, and ability of PI to pass through damaged membrane of dead cells and intercalate into nucleic acids. Therefore, this staining allows to distinguish between viable, early apoptotic, late apoptotic and necrotic cells. Incubations were performed as described above. Next, the organoids were dissociated in TrypLE Express (ThermoFisher), suspended in DPBS and centrifuged at 300g for 5 min. Resuspended pellets underwent AV-FITC/PI staining for FC analysis, per the manufacturer’s instructions. Cell suspensions were then analyzed by Beckman Coulter CytoFLEX. The obtained data were analyzed with use of Kaluza analysis 1.5A software (Beckman Coulter).

### 2.5 Protein expression analysis

Western blot (WB) was performed to evaluate the effect of the inhibitors on the level of α-syn (SNCA, pSer129) and ER stress markers (BiP, eIF2α/p-eIF2α, CHOP, XBP1, ATF6) in PD organoids. Incubations were performed as described above. Next, the organoids were lysed using Minute™ Total Protein Extraction Kit (Invent Biotechnologies) supplemented with protease and phosphatase inhibitor cocktail (Thermo Scientific). Proteins were quantified using Pierce™ BCA Protein Assay (Thermo Scientific). Equal amounts of proteins were loaded for each sample and electrophoresed on 10% polyacrylamide gel (Thermo Scientific). Following SDS-PAGE, proteins were transferred onto nitrocellulose membranes fixed for 30 min at RT in 4% PFA and 0.1% glutaraldehyde. Membranes were incubated with 5% BSA/milk solution diluted in TBST for 1 h, then with primary antibodies overnight at 4°C and peroxidase-conjugated secondary antibodies for 1 h at RT. Image acquisition and densitometry were performed with a chemiluminescent ChemiDoc Imaging System (Bio-Rad). The list of the applied antibodies is provided in Table 3 below.

**Table 3.** The full names and catalog numbers of the reagents used in the immunoblot experiments.

| Target | Antibody name | Dilution | Company | Catalog No. |
| --- | --- | --- | --- | --- |
| TH | Anti-Tyrosine Hydroxylase Antibody | 1:1000 | Sigma-Aldrich | AB152 |
| Tuj1 | β3-Tubulin (D71G9) XP® Rabbit mAb | 1:1000 | Cell Signaling Technology | 5568 |
| α-synuclein | alpha Synuclein Monoclonal Antibody (Syn 211) | 1:500 | Invitrogen | 32-8100 |
| pS129 α-synuclein | Phospho-α-Synuclein (Ser129) (D1R1R) Rabbit mAb | 1:500 | Cell Signaling Technology | 23706 |
| BiP | BiP (C50B12) Rabbit mAb | 1:1000 | Cell Signaling Technology | 3177 |
| p-eIF2α | Phospho-eIF2α (Ser51) antibody | 1:500 | Cell Signaling Technology | 9721 |
| eIF2α | eIF2α (D7D3) rabbit mAb | 1:1000 | Cell Signaling Technology | 5324 |
| CHOP | CHOP (L63F7) Mouse Monoclonal Antibody | 1:1000 | Cell Signaling Technology | 2895 |
| XBP1s | XBP-1s (D2C1F) rabbit mAb | 1:500 | Cell Signaling Technology | 12782 |
| ATF6 | ATF6 Polyclonal Antibody | 1:500 | Proteintech | 24169-1-AP |
| β-Actin | β-Actin (13E5) rabbit mAb | 1:1000 | Cell Signaling Technology | 4970 |
| Rabbit Ab | Anti-rabbit IgG, HRP-linked antibody | 1:5000 | Cell Signaling Technology | 7074 |
| Mouse Ab | Goat anti-Mouse IgG (H+L) Secondary Antibody, HRP | 1:5000 | Invitrogen | 31430 |

### 2.6 Gene expression analysis

The effect of the inhibitors on the expression of ER stress-associated genes in PD organoids was assessed by qRT-PCR analysis. Incubations were performed as described above. To isolate total RNA, the organoids were homogenized and lysed using PureLink RNA Mini Kit (Invitrogen) as described in the manufacturer’s instruction. The total RNA obtained was quantified by the Synergy HT Microplate Reader (BioTek). After conducting reverse transcription following the protocol of the High Capacity RNA to DNA Kit (Thermo Fisher Scientific), qRT-PCRs were performed using TaqMan Gene Expression Master Mix (Thermo Scientific). Amplification was performed in a Bio-Rad CFX96 detection system (Bio-Rad) as follows: an initial denaturation 15 min at 95°C, 40 cycles of denaturation for 15 s at 95°C, annealing for 30 s at 60°C, and elongation for 30 s at 72°C. The obtained data was analyzed using 2^–ΔΔCt^ method. The list of the applied TaqMan probes is provided below in Table 4.

**Table 4.** The catalog numbers of the applied TaqMan® Gene Expression Assays.

| Encoded protein | Target gene | Company | Assay ID |
| --- | --- | --- | --- |
| α-synuclein | SNCA | Applied Biosystems | Hs00240906_m1 |
| BiP | HSPA5 | Applied Biosystems | Hs99999174_m1 |
| eIF2α | EIF2A | Applied Biosystems | Hs00230684_m1 |
| CHOP | DDIT3 | Applied Biosystems | Hs01090850_m1 |
| XBP1 | XBP1 | Applied Biosystems | Hs00231936_m1 |
| ATF6 | ATF6 | Applied Biosystems | Hs00232586_m1 |
| β-actin | ACTB | Applied Biosystems | Hs99999903_m1 |

### 2.7 Statistics

Statistical analysis was performed by STATISTICA 13.3 software (StatSoft). Significance was evaluated with a two-tailed unpaired t-test (for normally distributed data) or a two-tailed Mann–Whitney U test (for not normally distributed data) when two conditions were compared. For more than two groups comparison, an ANOVA test followed by a Tukey’s post hoc test (for normally distributed data) or a Kruskal–Wallis test followed by a Dunn’s post hoc test (for not normally distributed data) were performed. P-values<0.05 were considered significant.

## 3. Results

### 3.1 The morphological features of midbrain organoids

We observed the presence of EBs as soon as 1 day after seeding of iPSC into microwells. The EBs were maintained in the microwells with daily media change to ensure the proper differentiation towards midbrain fate and organoid tissue formation. After day 7, when the preformed organoids were visible to the naked eye, they were transferred to the suspension stationary culture in the ultra-low attachment plates with concomitant media composition change to enable organoid expansion. The progressive growth of tissue in these conditions was apparent by multiple bud extrusions, which gave the organoid flower-like shape. Once the culture reached day 25, the organoids were transferred into differentiation medium – at this point the tissue growth became less rapid and the initially spherical microtissues developed more irregular shape, typical for organoid. At day 43, the midbrain differentiation process was completed and the organoids were from now on grown in the maintenance medium. After 55 days of culture, the organoids had approximately two and half millimeters in diameter (Fig. 1).

**Fig. 1.**
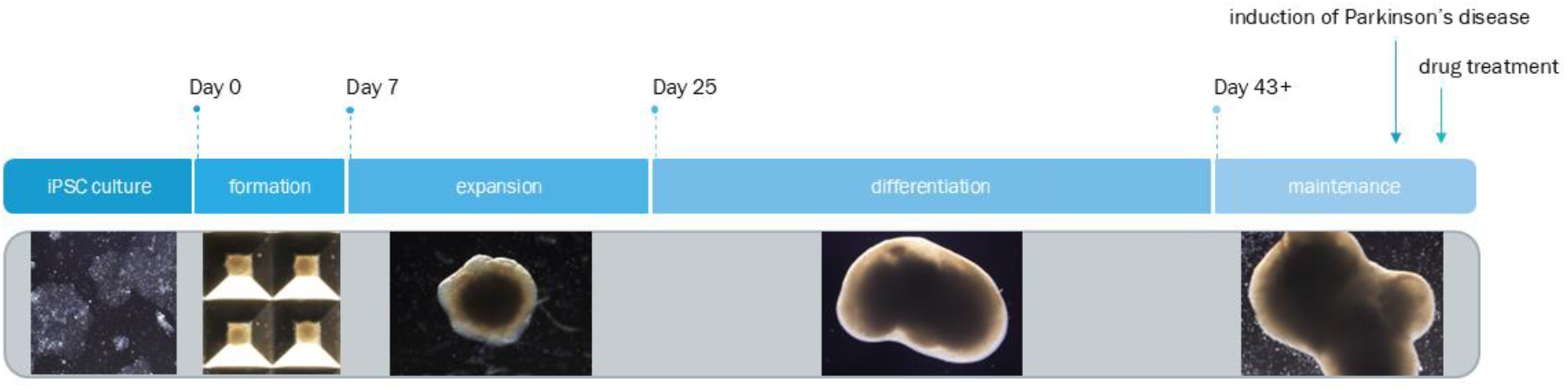
A schematic diagram representing generation of human midbrain organoids from an iPSC line obtained from a healthy donor. Following the final differentiation of the organoids, pathological Parkinson’s disease-specific conditions were induced via incubation with α-synuclein performed fibrils and 6-hydroxydopamine. After induction of neurodegeneration (48 h), the drug treatment was performed using anti-aggregation agent anle138b and PERK inhibitor AMG44.

The hematoxylin/eosin staining of the 55-day organoids showed a typical cytoarchitecture of brain tissue with multiple rosettes. The organoids consisted of the outer layer of proliferating cells (neurotrophic zone/organoid rim), the middle layer of differentiated cells (maturation zone/organoid core) and the inner layer consisting of accumulated dead cells (hypoxia zone/necrotic core) (Fig. 2).

**Fig. 2.**
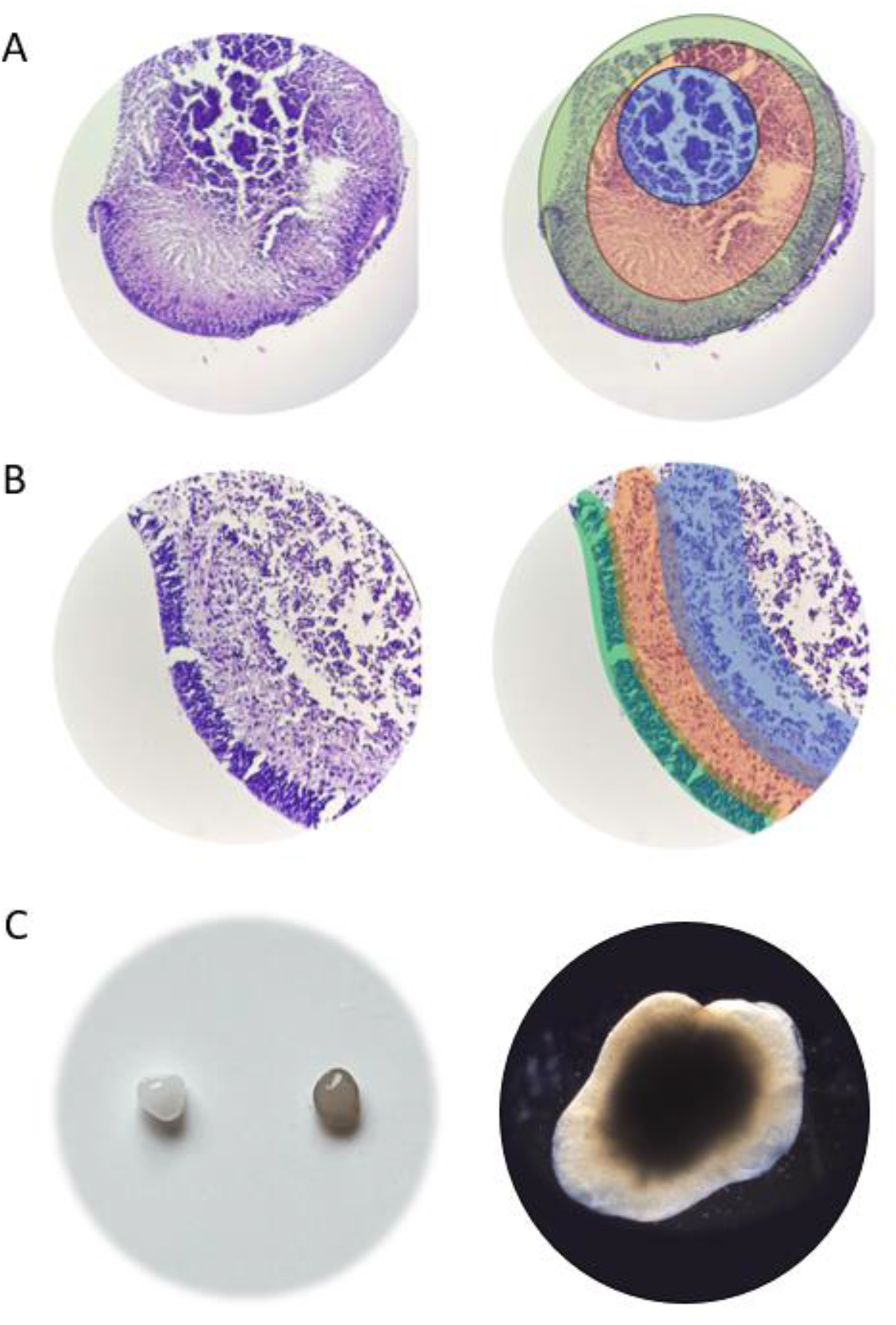
The histological examination of tissue architecture in 55-day-old human midbrain organoids using typical hematoxylin and eosin staining (A,B). The color lining (A; right) shows organization of the organoid structure: outer layer of organoid rim (green), middle layer of organoid core (orange), and inner layer of necrotic core (blue); magnification 40X. The color marking of the organoid margin (B; right) outlines the outer neurotrophic zone (green), middle maturation zone (orange) and inner hypoxia zone (blue); magnification 100X. Macroscopic image (C; left) of human midbrain organoid (left) and dopamine-stimulated human midbrain organoid (right) reveal color changes as an effect of dopamine exposure and the resulting neuromelanin accumulation. Microscopic image (C; right) shows irregular shape and dense structure typical for the organoid; magnification 40X.

The presence of neuronal and dopaminergic markers were assessed by immunofluorescence and WB. Immunostainings of 30-day organoids revealed abundance of FOXA2 in the center of the rosettes containing neuronal progenitor cells. At day 55, we observed an enrichment of TH-positive dopaminergic neuronal cells, high rate of neurons expressing MAP2 and NFP, and a number of glial cells positive for S100β, typical for differentiated midbrain organoid (Fig. 3).

**Fig. 3.**
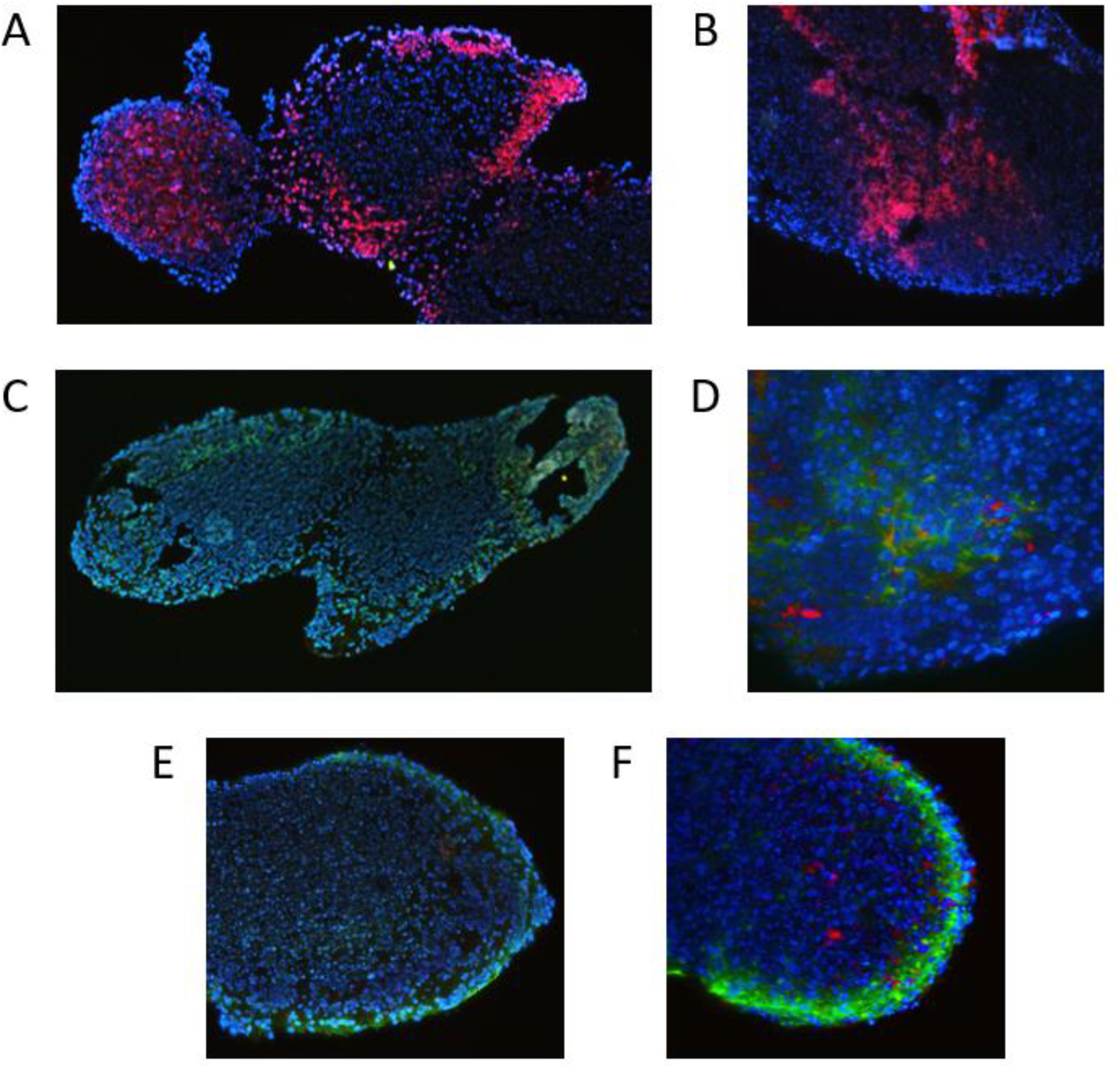
The representative visualization of differentiation markers in human midbrain organoids detected by immunofluorescence (A-F). Organoids at day 30 were strongly positive for FOXA2 (red signal), indicating the zones containing neural progenitors (A,B). Mature 55-day organoids exhibited elevated expression of neuronal marker MAP2 (green signal; C,D), dopaminergic marker TH (red signal; C,D) axon marker NFP (green signal; E,F) and glial cell marker S100β (red signal; E,F). Hoechst was used for counterstaining. Images were taken at magnifications 100X (A,C,E), 200X (B,F), and 400X (D).

A separate immunoblot experiment showed that the organoid lysates were characterized by the expression of dopaminergic marker TH, neuronal marker Tuj1, and α-synuclein compared to iPSC line sample (Fig. 4).

**Fig. 4.**
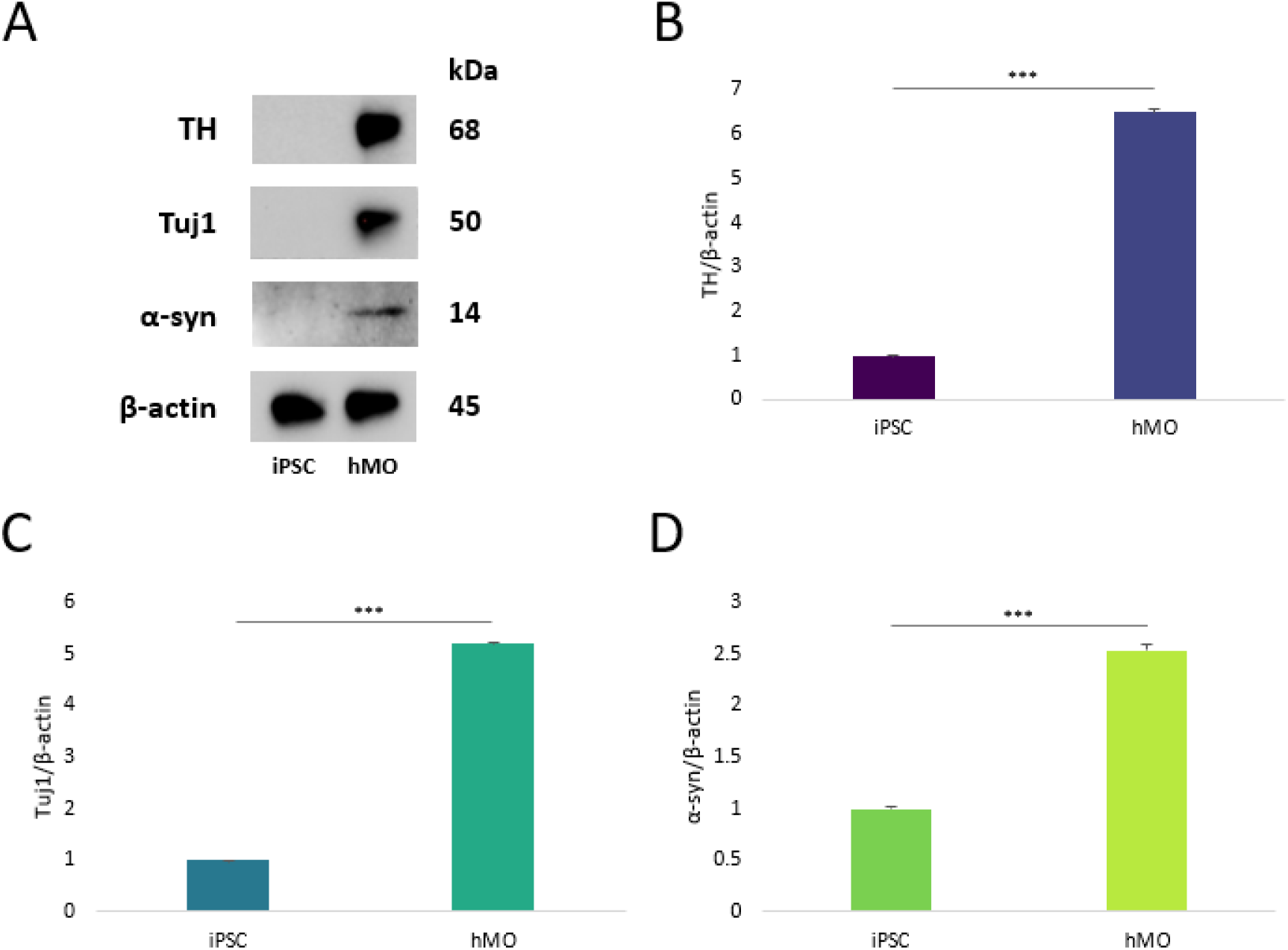
The Western blot analysis demonstrates a significant increase in the expression of dopaminergic neuronal marker tyrosine hydroxylase (TH), neuronal marker β tubulin III/Tuj1, and synaptic protein α-synuclein (α-syn) after differentiation of induced pluripotent stem cells into human midbrain organoids. The figure shows immunoblot images (A) and normalized protein levels for TH (B), Tuj1 (C) and α-syn (D). β-actin was used as a loading control. The Student’s t-test was used in the statistical analysis. Data are expressed as mean ± SD. *** p < 0.001 vs. iPSC. Abbreviations: iPSC – induced pluripotent stem cells; hMO – human midbrain organoids.

### 3.2 The confirmation of amyloid formation by α-syn pre-formed fibrils

The thioflavin T emission curves demonstrate increased fluorescence in response to α-syn aggregation and formation of β-sheet-rich structures. While the thioflavin T dye alone and α-syn monomer showed no fluorescent signal, α-syn PFFs induced substantial increase of fluorescence intensity over time. The thioflavin T-binding reaction kinetics rapidly increased upon incubation of α-syn PFFs combined with α-syn monomer, compared to PFFs alone and monomer alone. The assay confirmed the ability of applied α-syn PFFs to seed the formation of new α-syn fibrils from the pool of α-syn monomers (Fig. 5).

**Fig. 5.**
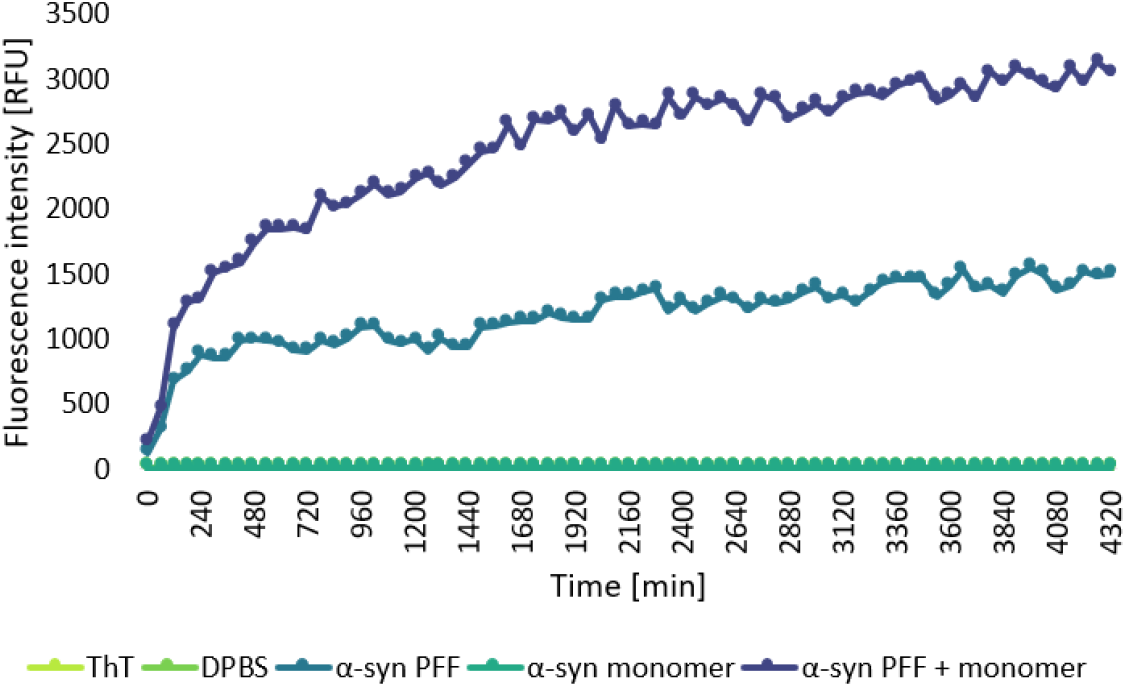
The thioflavin T (ThT) assay confirms the aggregation of α-synuclein (α-syn). The kinetic curves represent the concentration-dependent increase in ThT fluorescence intensity induced by aggregated α-syn species (preformed fibrils/PFFs, PFFs + monomer).

### 3.3 Treatment of PD organoids with anle138b and AMG44 increases cell viability

The cell viability of midbrain organoids was assessed by the specialized assay adjusted to 3D culture conditions. The analysis demonstrated that the small-molecule inhibitors anle138b and AMG44 show no substantial cytotoxicity towards midbrain organoids at any tested concentration as compared to vehicle control (Fig. 6 A,B). In the next step of the cell viability experiment, we assessed the effect of each inhibitor alone on the viability of PD organoids at 0.1-200 µM and at 24 h post-treatment. Treatment of PD organoids with AMG44 as well as anle138b resulted in a significant increase in cell viability as evidenced by higher ATP content (Fig. 6 C,D). These results are in accordance with the previous findings, which demonstrated that AMG44 is effective against neurotoxic damage in vitro [33], and that anle138b effectively rescues neurodegeneration after the onset of pathogenic α-syn accumulation in vivo [26]. Finally, the combination of both drugs at their effective concentrations (25 µM AMG44 and 25 µM anle138b at 24 h post-treatment) was evaluated in PD organoids. Strikingly, application of both drugs led to a significant increase in the number of metabolically active cells within the organoid as compared to treatment with each inhibitor alone (Fig. 6 E).

**Fig. 6.**
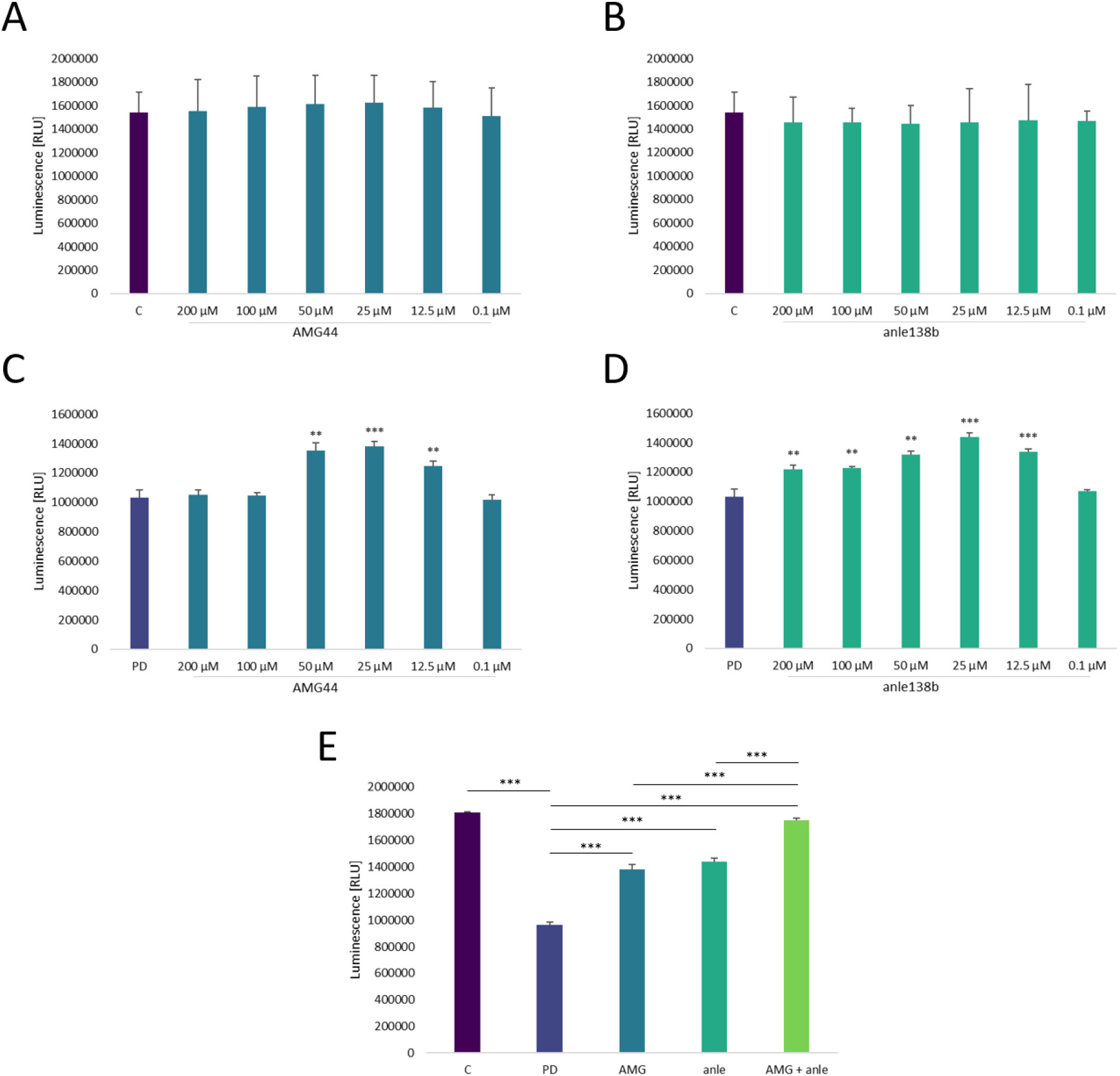
Cytotoxicity analysis of PERK inhibitor AMG44 (A) and α-synuclein aggregation inhibitor anle138b (B) in midbrain organoids, and the effect of post-treatment with AMG 44 (C), post-treatment with anle138b (D) and combined treatment with AMG44 and anle138b (E) in midbrain organoids with induced Parkinson’s disease (PD). T-Student’s test (A-D) and one-way ANOVA (E) were used for the statistical analysis. Data are expressed as mean ± SD (n=3). **p<0.01, ***p<0.01 for C (A-B), PD (C-D) and all groups (E). Abbreviations: C – control organoids; PD – organoids with induced PD; AMG – PD organoids treated with AMG44 inhibitor; anle138b – PD organoids treated with anle138b inhibitor; AMG+anle – PD organoids treated with AMG44 and anle138b inhibitors.

### 3.4 anle138b and AMG44 treatment reduces α-syn aggregation in PD organoids

The anti-aggregation effect of the inhibitors was evaluated by immunofluorescence staining, in which the presence of several species of α-syn was assessed (aggregated/plaques, p-Ser129 and total α-syn). PD organoids showed significant increase in fluorescent signal corresponding to aggregated and Ser129-phosphorylated α-syn in the organoid core and rim, with concomitant decrease in total α-syn level compared to control. Application of anle138b or AMG44 led to partial reduction of plaques and pSer129 intensity in PD organoids, with the levels of total α-syn signal remaining unaffected. Treatment with the combination of both drugs was able to significantly decrease the rate of α-syn aggregation and Ser129 phosphorylation, and restore the total α-syn intensity, indicative of higher amount of unaggregated α-syn forms within the sample. This effect was comparable to the levels of the respective forms of α-syn in the control organoid (Fig. 7).

**Fig. 7.**
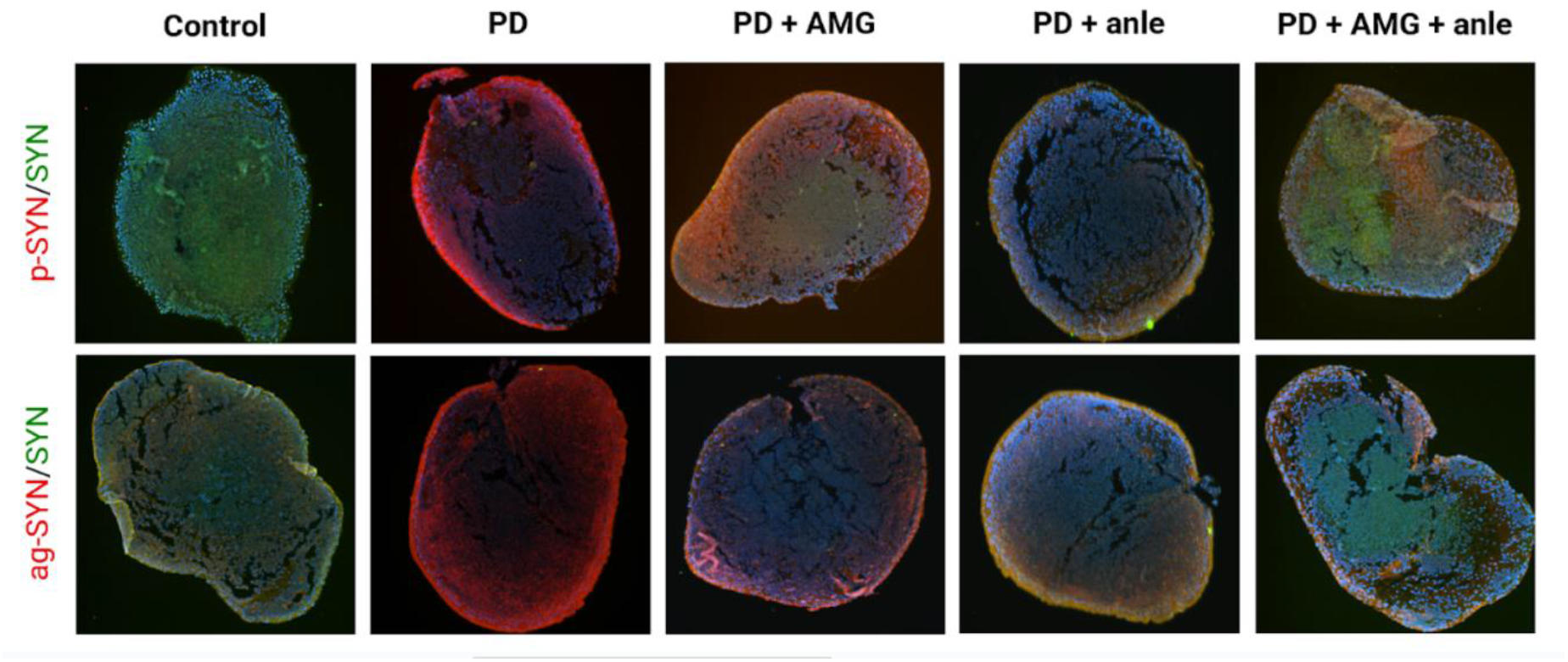
Representative images from immunofluorescence staining of the pathological forms of α-synuclein (α-syn): Ser129-phosphorylated α-syn (red signal; top row) and α-syn aggregates (red signal; bottom row) together with total α-syn (green signal) and nuclear counterstaining (Hoechst; blue signal) in the paraffin-embedded sections of normal and PD midbrain organoids treated with anle138b and AMG44 inhibitors. Abbreviations: PD – organoids with induced Parkinson’s disease (PD); AMG – PD organoids treated with AMG44 inhibitor; anle138b – PD organoids treated with anle138b inhibitor; AMG+anle – PD organoids treated with AMG44 and anle138b inhibitors.

### 3.5 Application of anle138b and AMG44 in PD organoids decrease the apoptosis rate

FC analysis using FITC-AV/PI fluorescent staining revealed relatively high rate of cell death within the control unaffected organoids (which is acceptable giving the presence of large necrotic cores in the mature ∼60-day organoids [41]). We observed significantly increased rate of apoptosis in PD organoids as compared to control. Upon single treatment of PD organoids with anle138b or AMG44 inhibitor, the apoptotic cell rate slightly decreased, but the difference was statistically insignificant. Incubation of PD organoids with both inhibitory compounds resulted in a significant decrease in the number of apoptotic cells as compared to PD group and the groups treated with each inhibitor in monotherapy (Fig. 8).

**Fig. 8.**
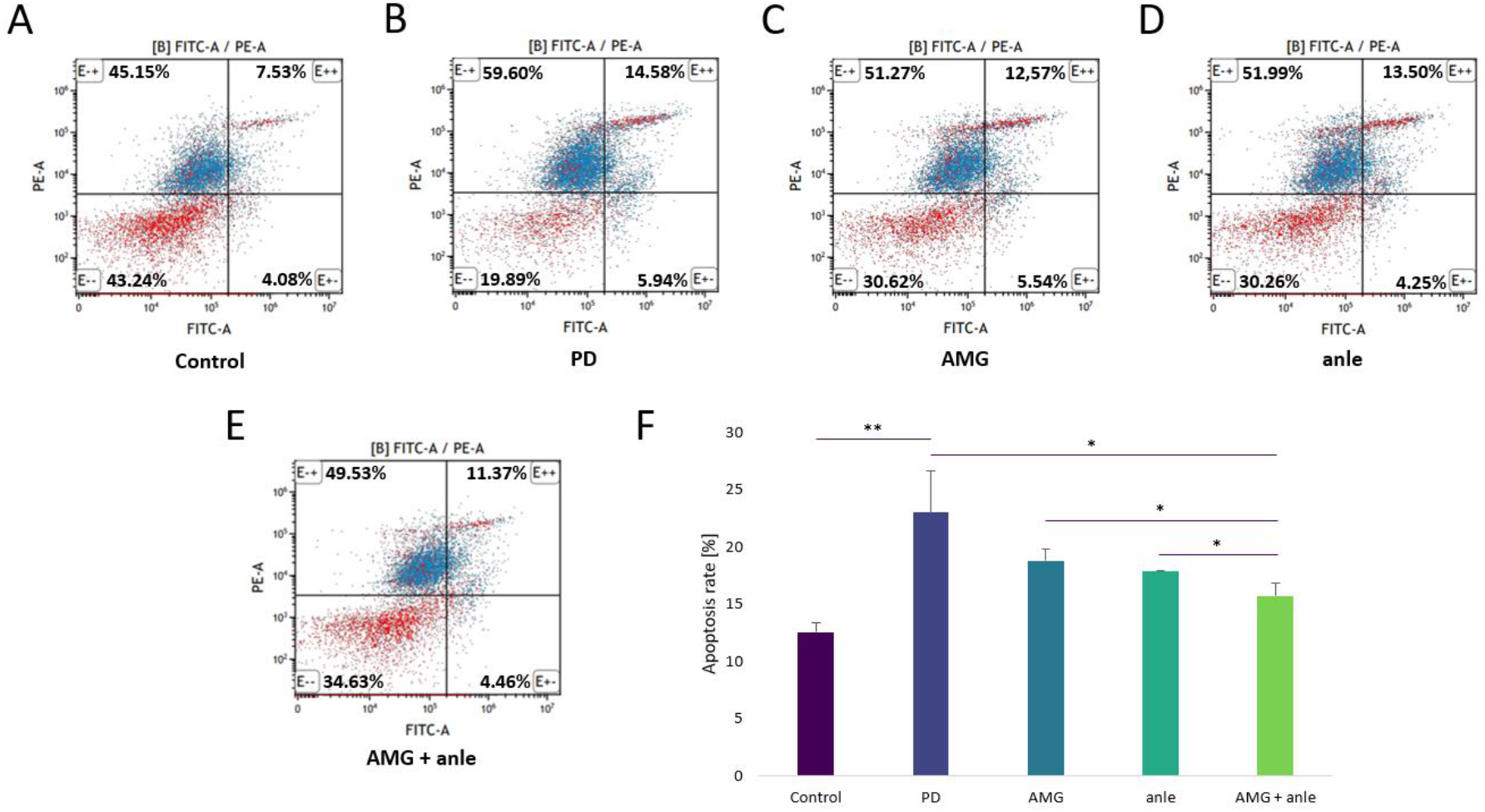
Flow cytometric Annexin V-FITC/Propidium iodide (AV/PI) dual staining analysis of apoptosis in PD organoids treated with anle138b and AMG44 inhibitors. Dot plot graphs illustrate the percentage of viable cells (lower left Q3 quadrants; AV-/PI-), early apoptotic cells (lower right Q4 quadrants; AV+/PI-) late apoptotic cells (upper right Q2 quadrants; AV+/PI+) and necrotic cells (upper left Q1 quadrants; AV-/PI+) cells (A-E). Bar graph shows quantified percentage of apoptotic scells (F). One-way ANOVA was used for the statistical analysis. The data are expressed as mean ± SD (n=3). *p< 0.05, **p < 0.01 for all groups. Abbreviations: C – control organoids; PD – organoids with induced Parkinson’s disease (PD); AMG – PD organoids treated with AMG44 inhibitor; anle138b – PD organoids treated with anle138b inhibitor; AMG+anle – PD organoids treated with AMG44 and anle138b inhibitors.

### 3.6 anle138b and AMG44 rescue PD neurodegeneration by modulating the expression of α-syn and ER stress markers

PD organoids expressed significantly higher levels of total α-syn, Ser129-phosphorylated α-syn monomers and fibrils in comparison with normal organoids. Treatment with AMG44 and anle138b significantly reduced the total and pSer129 α-syn levels, with anle138b exhibiting slightly stronger anti-aggregation effect (which results from its direct mechanism of action). Combination of both compounds more effectively reduced the rate of α-syn phosphorylation at Ser129 than each inhibitor alone. Of note, none of the compounds completely abolished α-syn expression, which is important for the maintenance of physiological synaptic function (Fig. 9).

**Fig. 9.**
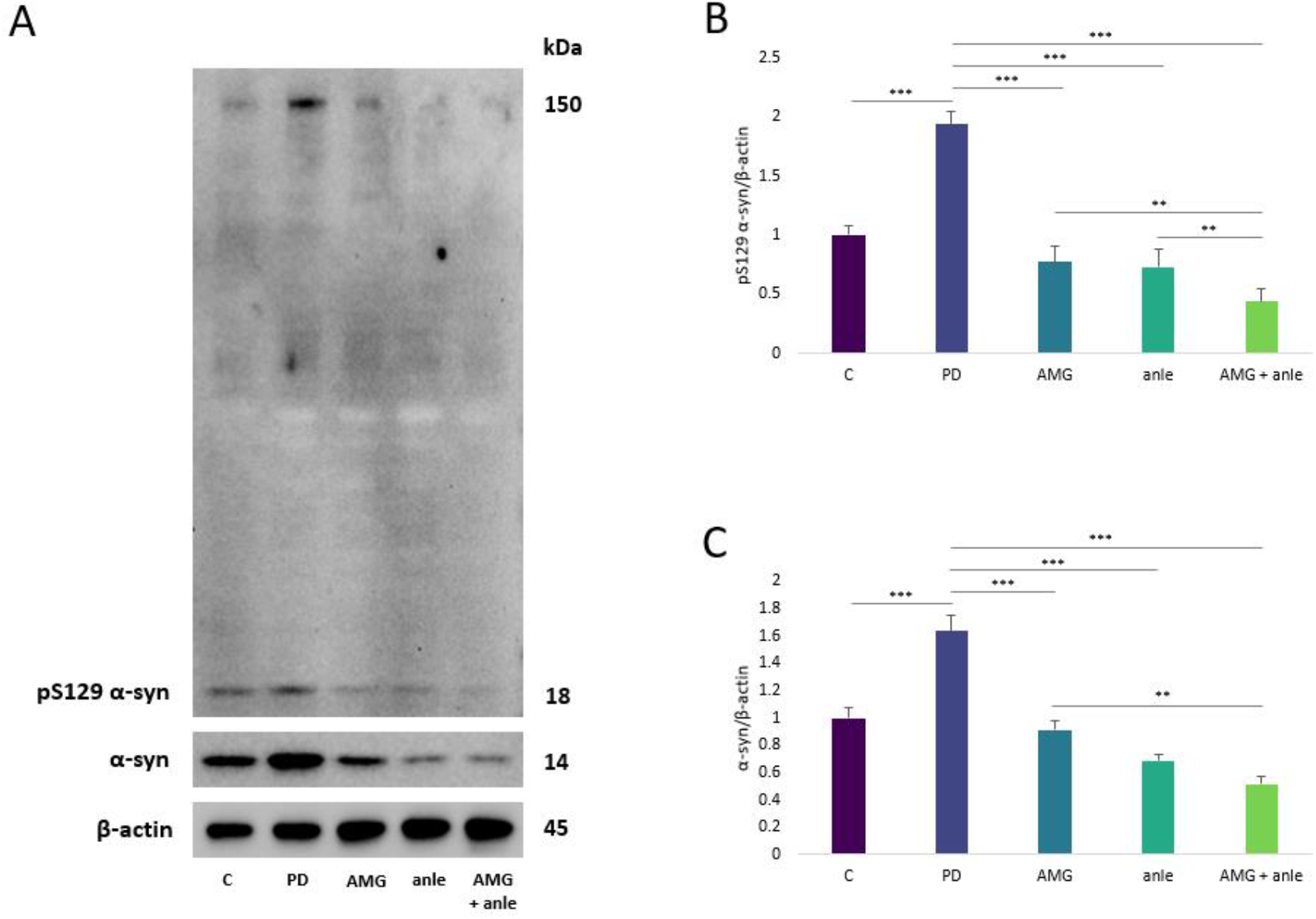
The effect of anle138b and AMG44 inhibitors on the expression of total and S129-phosphorylated α-synuclein (α-syn) in Parkinson’s disease (PD) midbrain organoids evaluated by immunoblotting. The figure depicts Western blot images (A) and normalized protein expression levels of pS129 α-syn (B) and total α-syn (C). Abbreviations: C – control organoids; PD – organoids with induced PD; AMG – PD organoids treated with AMG44 inhibitor; anle138b – PD organoids treated with anle138b inhibitor; AMG+anle – PD organoids treated with AMG44 and anle138b inhibitors.

The expression level of chaperone BiP protein was decreased in PD organoids compared to control, and it was effectively restored only by combination of anle138b and AMG44. PD organoids showed increase in eIF2α phosphorylation, and the p-eIF2α/eIF2α ratio was effectively reduced by each inhibitor. Importantly, the inhibitors treatment did not completely suppress eIF2α phosphorylation, which could lead to detrimental effects. The level of other PERK downstream effector, CHOP, was significantly upregulated in PD organoids, and it was substantially downregulated by the inhibitors treatment. In comparison with single treatments, the combination of anle128b and AMG44 significantly upregulated the proteins XBP1s and ATF6, which are known to play a cytoprotective role in the course of PD (Fig. 10).

**Fig. 10.**
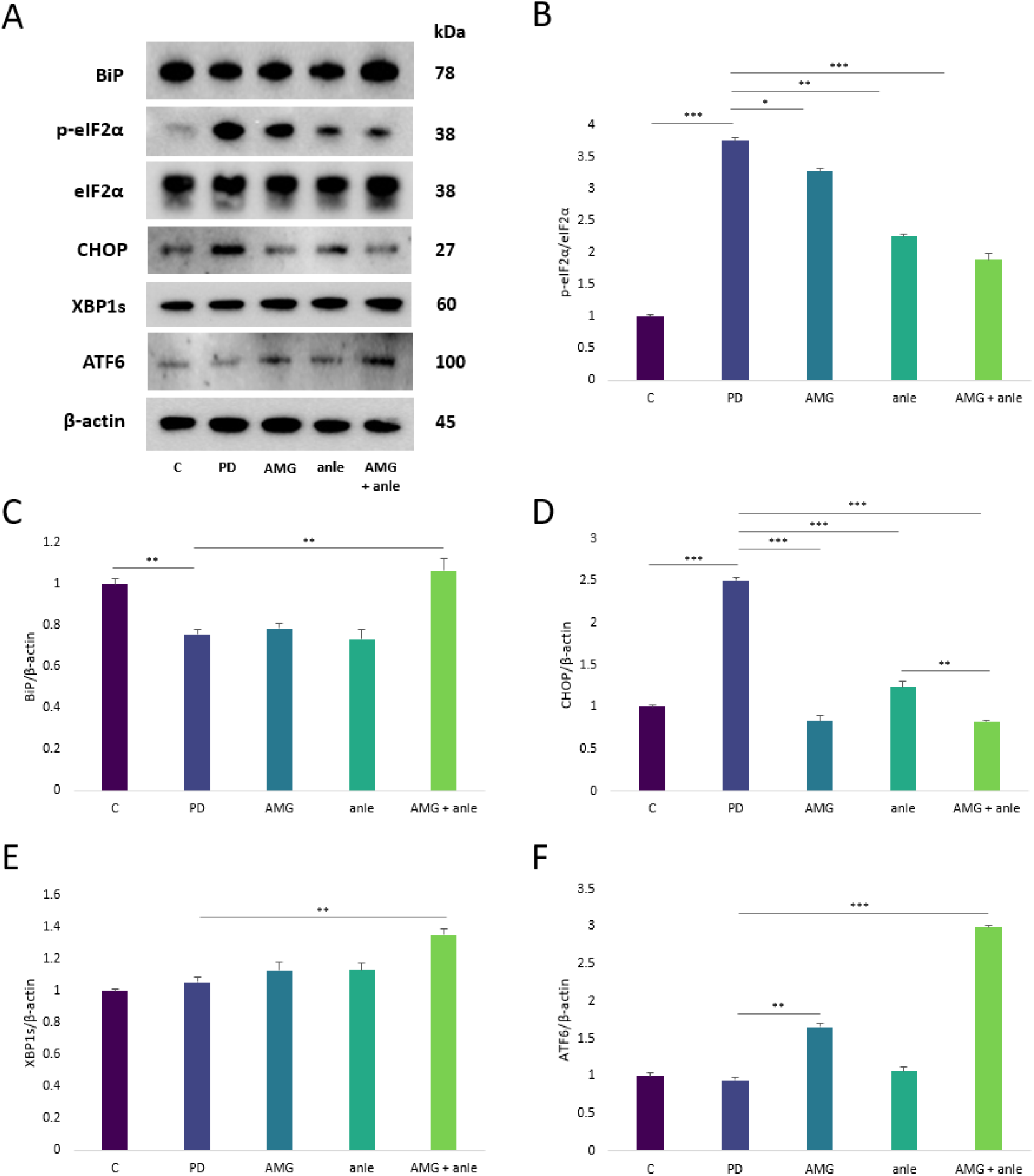
The Western blot images (A) and normalized protein expression levels of the endoplasmic reticulum stress-associated factors (B–F): p-eIF2α (B), BiP (C), CHOP (D), XBP1s (E), and ATF6 (F) in Parkinson’s disease (PD) midbrain organoids treated with anle138b and AMG44 inhibitors. β-actin was used as a loading control. One-way ANOVA was used for the statistical analysis. The data are expressed as mean ± SD (n=3). *p< 0.05, **p < 0.01, ***p < 0.001 for all groups. Abbreviations: C – control organoids; PD – organoids with induced PD; AMG – PD organoids treated with AMG44 inhibitor; anle138b – PD organoids treated with anle138b inhibitor; AMG+anle – PD organoids treated with AMG44 and anle138b inhibitors.

### 3.7 anle138b and AMG44 compounds act differently on the expression of α-syn- and ER stress-related genes

The gene expression analysis revealed a significant increase in the level of *SNCA* mRNA encoding α-syn as well as all evaluated UPR-associated genes (*HSPA5, EIF2A, DDIT3, XBP1* and *ATF6*) in PD organoids compared to control. Interestingly, treatment with each inhibitor exerted different effect on *SNCA* expression –while AMG44 strongly downregulated *SNCA* mRNA, anle138b significantly upregulated it, and this could be explained by various mechanisms of action of the drugs. Furthermore, while single treatment with AMG44 had negligible effect or resulted in a significant decrease in the expression of ER stress-related genes (which is an expected effect for PERK inhibitor), anle138b acted mainly by upregulating the level of these mRNA; this suggests that anle138b anti-aggregation treatment may evoke an early ER stress response at gene expression level. Combination of both substances resulted in normalization of *SNCA* mRNA level and led to a significant increase in the expression of *HSPA5, EIF2A, XBP1* and *ATF6* mRNA compared to single treatments. Therefore, it appeared that combined treatment with AMG44 and anle138b upregulated GRP78 chaperone and protective branches of the UPR related to IRE1- and ATF6-depenent signaling at the mRNA level. Moreover, addition of AMG44 significantly decreased the expression of pro-apoptotic *DDIT3* gene compared to single anle138b treatment, suggesting that AMG4 alleviated anle138b-induced terminal UPR activation (Fig. 11).

**Fig. 11.**
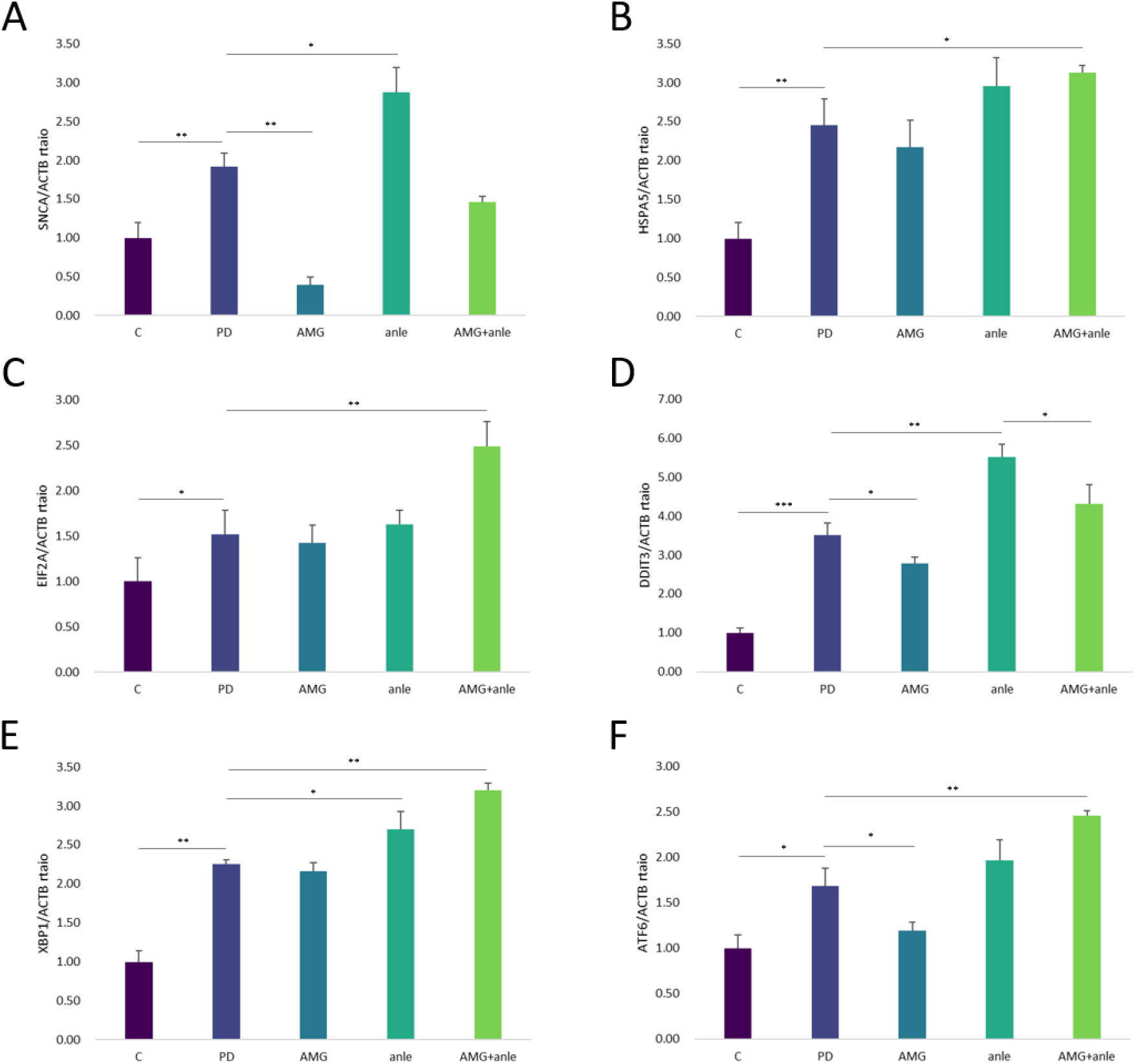
The mRNA expression levels of *SNCA* gene encoding α-syn (A) and endoplasmic reticulum (ER) stress-related genes *HSPA5* (B), *EIF2A* (C), *DDIT3* (D), *XBP1* (E), *ATF6* (F) in PD organoids treated with anle138b and AMG44 inhibitors. *ACTB* was chosen as a reference gene. One-way ANOVA was used for the statistical analysis. Values are presented as mean ± SD (n = 3), *p < 0.05, **p < 0.01, ***p < 0.001 for all groups. Abbreviations: C – control organoids; PD – organoids with induced Parkinson’s disease (PD); AMG – PD organoids treated with AMG44 inhibitor; anle138b –PD organoids treated with anle138b inhibitor; AMG+anle – PD organoids treated with AMG44 and anle138b inhibitors.

## 4. Discussion

In the present study, we evaluated for the first time the potential neuroprotective properties of the small-molecule substances anle138b and AMG44 both separately and in combination in the novel human-based, in vitro 3D model of sporadic PD. The obtained results show that treatment of PD organoids with the investigated compounds results in a significant increase in cell metabolic activity, reduction of neuronal cell death, and decrease in accumulation of the two independent pathogenic forms of α-syn, namely aggregated α-syn and pS129 α-syn. In each case, application of both drugs was significantly more effective at providing neuroprotection than treatment with each inhibitor alone. At the mechanistic level, treatment with the compounds significantly reduced the levels of total and pSer129 α-syn and modulated the expression of ER stress markers. Combination of both inhibitors was more effective a decreasing the rate of α-syn Ser129 phosphorylation than each substance alone. Moreover, only combination of anle138b and AMG44 effectively upregulated the BiP, XBP1s and ATF6 factors at protein and mRNA level, which suggests switching UPR towards its adaptive, pro-survival arm. All mentioned UPR factors had previously been reported to have a protective role in PD pathogenesis by numerous independent studies [15, 42–46]. We also for the first time demonstrate a new 3D model of sporadic PD, which represents the most important features of human pathology – reduced metabolic activity, the presence of specific α-syn aggregates and α-syn phosphorylated at Ser129, and increased rate of apoptosis.

The rationale behind combining anti-aggregation therapy with inhibition of ER stress lies in a direct mechanistic link between protein misfolding and cellular stress. Accumulation of misfolded α-syn in the form of toxic oligomers or aggregates disrupts cellular proteostasis, resulting in ER stress, UPR activation and apoptotic cell death [12, 36]. While anti-aggregation approaches reduce the formation and toxicity of pathogenic α-syn species, they do not directly alleviate the ER stress response, restore the proteostasis or prevent UPR-mediated apoptosis in already stressed neuronal cells. On the other hand, ER stress modulators such as PERK inhibitors can restore the translational control and limit ER-stress-induced apoptosis, but may not directly affect the ongoing aggregation. Combining both approaches addresses the two pathogenic events associated with α-syn toxicity (aggregates formation and ER stress), potentially enhancing neuroprotection, preserving neuronal function, and slowing disease progression more effectively than either strategy alone. We speculate that addition of PERK inhibitor to the anti-aggregation agent may help facilitate the cleanup of accumulated α-syn [47]. Of note, the expression of such proteins as α-syn or p-eIF2α was substantially reduced but not completely silenced by the inhibitor treatment. This is important in view of the fact that these factors are key in maintaining essential physiological functioning of neuronal cell, and lack of their normal activity might carry devastating consequences [48, 49]. Partial inhibition of a target protein is a desirable drug feature as it enables preservation of vital cellular processes while reducing pathological activity [32]. Such partial or gradual target blockade can also more closely mimic physiological regulation and lower the risk of adverse effects; this is especially beneficial in chronic diseases like PD which require prolonged treatment.

We previously evaluated the neuroprotective properties of AMG44 and JNK V inhibitor in the neurotoxin-based 2D cellular models of PD [33, 50]. We found that in 6-OHDA model, JNK inhibition exerted stronger neuroprotective effect than PERK inhibition, whereas in a separate study both inhibitors seemed to be equally effective against rotenone-induced neurodegeneration. We also identified the main difference in the mechanism of action of both drugs – PERK inhibitor acts mainly by decreasing caspase-3 level and apoptosis, whilst JNK inhibitor is a more potent inhibitor of necrosis. We speculate that the discrepancies between the two models could be explained by different mechanisms of toxicity induced by the two neurotoxins, and they could also have resulted from the applied model (retinoic acid-differentiated SH-SY5Y neuroblastoma cell line), which, although widely used as an in vitro model of PD, possesses several limitations. As these cells are a transformed cancer cell line, even after differentiation they do not fully acquire the molecular properties of mature human dopaminergic neurons, and their differentiation process is heterogeneous and highly sensitive to protocol variables [51]. Also, these cells do not normally express high levels of α-syn, a key protein in PD pathogenesis (unless it is chemically or genetically induced [52]), and the neurotoxin treatment alone seems to be a more suitable model for studying acute toxicity rather than chronic neurodegenerative process. Although our previous results were in favor for JNK inhibition, literature data suggest that these enzymes possess many important physiological functions [53], so suppression of endogenous JNK in the CNS might be not quite favorable [54]. Despite exhibiting disease-modifying effects in vivo, mixed lineage kinase inhibitor CEP-1347 which is also a JNK1 inhibitor failed to affect disability in early PD patients in a clinical trial [55]. Some studies also suggest that pathological α-syn may in fact decrease neuronal JNK activity – in such case, further inhibition of JNK seems to be pointless [56, 57]. We have also recently found that higher concentrations of JNK V inhibitor are in fact cytotoxic towards undifferentiated SH-SY5Y neuroblastoma cells, so it could have a better potential as an antineoplastic agent than neuroprotector [58]. Considering the above and given the uncertain status of AMG44 effectiveness against PD based on previous experiments results, it was selected for further testing in a 3D model to fully validate its neuroprotective potential in more specific PD conditions.

The present findings are consistent with literature reports of a potent anti-aggregation effect of anle138b against PD. Anle138b showed wide neuroprotective effects in previous studies, which include reduction of amyloid proteins aggregation, preservation of neuronal function, and improvement of disease-related phenotypes. It was extensively studied in multiple preclinical models of neurodegenerative diseases like PD, AD, MSA and prion disease with favorable outcomes [26–28]. Its pleiotropic mechanism results from the fact that anle138 not only targets pathological misfolding and toxic oligomer formation of α-syn, but also prion protein and tau [26, 59]. Simultaneously, it does not substantially affect α-syn monomers [26], which suggests a specificity for the pathogenic aggregated species. Besides broad anti-aggregation activity against proteins relevant to multiple neurodegenerative disorders, the compound is characterized by good brain penetration and oral bioavailability [26]. In contrast to currently used therapies for PD, anle138b offers a potential disease-modifying strategy that may stop the disease progression rather than provide purely symptomatic relief. Nevertheless, given its broad anti-aggregation effect, the target specificity of the compound is not fully resolved, which raises concerns about potential off-target events. Additionally, there is uncertainty regarding long-term safety of the compound, optimal dosing strategies, and clinical efficacy in humans, all of which require thorough validation in advanced clinical trials. Moreover, in MSA mouse model, treatment with anle138b only partially improved motor function without preventing neuronal loss and neuropathological changes in the brain [60], whereas in the mouse models of inherited prion disease, anle138b therapy did not significantly extend disease-free survival or affect accumulation of mutant prion protein [61]. It seems therefore that the above-mentioned pathologies might have been too advanced to benefit from anle138b treatment, and that the efficacy of anle138b depends on disease stage and aggregation burden. In general, a major challenge of anti-aggregation therapies in neurodegeneration is that protein aggregates occur at later disease stages, when the neuronal damage is already extensive, and their ability to reverse pathology is limited. Notably, neurodegenerative diseases are often characterized by multifaceted pathologies, of which aggregation represents only one aspect – other molecular events like chronic ER stress, neuroinflammation, mitochondrial dysfunction, and synaptic loss may continue despite clearance of protein aggregates, which reduces the overall therapeutic effect [62–64]. It was also suggested that disrupting α-syn fibrils into smaller species without effective clearance may transiently increase the toxic oligomer populations, which are potent ER stress inducers [34, 35]. Therefore, combination of anti-aggregation agents with targeted immunotherapy, neuroprotective agents, or other disease-modifying treatments may amplify its benefits. Consistently, it was previously shown that combined treatment with anle138b and AFFITOPE® PD03 resulted in reduced IgG binding in the brains of MSA mice, suggesting lower burden of α-syn oligomers in the central nervous system [65]. Such multi-pronged approaches might help overcome the limitations of anti-aggregation therapies and improve potential clinical outcomes.

PERK inhibitors have been extensively explored in PD as a strategy to modulate chronic ER stress and the maladaptive UPR, which are implicated in α-synuclein aggregation, neuroinflammation and dopaminergic neuron loss, in particular when ER stress is sustained and pathological. In our previous works, we evaluated the potential neuroprotective effects of LDN-87357 and AMG44 PERK inhibitors in SH-SY5Y cellular models of PD, in which PERK inhibition resulted in, among others, increased cell metabolic activity, diminished apoptosis and modulation of ER stress markers [33, 66]. In a mouse PD model, oral administration of PERK inhibitor GSK2606414 protected dopaminergic neurons, reduced ER stress, restored synthesis of dopamine and synaptic proteins, and improved motor function [21]. However, as PERK signaling also plays essential role in maintaining cellular homeostasis, especially in neurons and secretory cells, systemic or prolonged PERK inhibition may exert significant toxicity (such as pancreatic dysfunction induced by GSK2606414 treatment [21]). Several off-target effects (e.g., inhibition of other kinases like RIPK1) of old-generation GSK inhibitors have also been reported [30], which also limits their potential clinical translation. It is therefore essential to develop new-generation PERK inhibitors with higher selectivity and limited adverse effects. Current efforts focus on achieving partial, transient, or brain-targeted PERK pathway modulation, or acting downstream of PERK (e.g. on eIF2α or CHOP) to balance neuroprotection in PD with maintaining safety. In line with this statement, an eIF2α phosphorylation modulator salubrinal protected SH-SY5Y cells against rotenone-induced ER stress and apoptosis [67], and it also decreased motor impairment and ameliorated neuroinflammation in a rat PD model [68]. To the best of our knowledge, the effect of UPR inhibitors have not yet been investigated against PD in human organoids. We also for the first time demonstrate potential anti-aggregation effect of the PERK inhibitor AMG44 in organoid PD model, a feature that has not been previously investigated with regard to UPR inhibitors. We hypothesize that modulation of ER stress by PERK inhibition may facilitate the degradation of α-syn aggregates via ER-associated degradation (ERAD) and ER-phagy pathway [69, 70], but further research is required to confirm this theory. Initial preclinical data suggest that AMG44 does not impair the pancreas function [31], but little is known regarding its oral bioavailability, pharmacokinetic and pharmacodynamic profile, and ability to cross blood-brain barrier; these aspects should be evaluated by future studies. Of note, in the present study, the effect of the drugs was investigated only at a single timepoint, and longer incubation periods which could correspond to chronic treatment have not been examined. Given the time-dependent nature of the UPR activity, we speculated that longer and more extensive anti-aggregation therapy could further aggravate ER stress and lead to stronger UPR activation (which was found by this study in particular at gene expression level). Thus, it would be beneficial to perform longer incubation periods and apply multiple drug concentration variants to fully explore the combination therapy effect.

Human midbrain organoids, although regarded as an excellent preclinical model of α-synucleinopathy [71], are not free of limitations. Most notably, the lack of vasculature of the organoid restricts oxygen and nutrient diffusion to its inner layers, leading to hypoxia and necrosis within the organoid core. Without vasculature it is also impossible to study blood-brain barrier integrity, which is an important aspect of PD pathogenesis [38]. The organoids may also show varying levels of maturity, miss some peripheral cell types and have limited functional connectivity compared to human midbrain [72]. The shortcomings related to vasculature and nutrient supply can be overcome by novel bioengineering strategies like co-culture with vascular cell types or vascular organoids, transplantation into rodent brains, use of microfluidic “organ-on-chip” systems, bioprinting or scaffold-based techniques [73]. Other improvements include extended culture times, refined differentiation protocols, and development of assembloids which better model physiological complexity of neural circuits [38]. Nonetheless, the effectiveness of the tested drugs in the organoid model provides strong evidence that their mechanisms of action are preserved in a more physiologically relevant context [71]. The next steps in this direction involves in vivo validation in appropriate PD animal models to assess their therapeutic efficacy under systemic conditions. This evaluation should also include safety and tolerability aspects, the effect on disease-relevant symptoms, as well as assessment of bioavailability, pharmacokinetics/pharmacodynamics, brain penetration, target engagement and optimal dosing. In vivo experiments (especially with regard to AMG44) should determine whether the observed 3D in vitro efficacy can translate into a viable therapeutic strategy in living organisms. Also, as the present study is limited by only one disease model and only one type of each inhibitor applied, our findings require further validation in preclinical studies. Future research should focus on evaluating other anti-aggregation drugs and UPR inhibitors in PD-specific conditions (especially genetic models) to better contextualize the pathological role of α-synucleinopathy-induced ER stress.

To date, many aggregation inhibitors for neurodegenerative diseases with proven in vivo efficacy failed to succeed in human clinical trials [74]. Such translational failure could be explained by fundamental differences between rodents and humans in terms of physiology, genetics, immune responses, metabolism and signaling pathways. Preclinical animal models often do not fully reflect the complexity, heterogeneity, and chronic nature of human disease, which results in overestimation of therapeutic benefit [39, 72]. A number of species-specific differences in drug metabolism, target engagement, and dose–response relationships may lead to insufficient efficacy or unexpected toxicity in humans [75]. Other variables include inadequate animal age, limited sample sizes, differences in molecular targets and preclinical/clinical endpoints, earlier time of intervention and relatively high drug doses applied to animals, all of which may affect the translation of preclinical results into human trials [76]. We believe that application of human organoids may facilitate the translation of preclinical findings into clinical trials as they provide a more physiologically relevant human in vitro system. Preclinical use of organoid cultures bridges the gap between conventional 2D cell culture and animal studies, which otherwise frequently fail to predict clinical efficacy [72]. The organoids represent key aspects of cellular diversity, human tissue architecture, and disease-relevant molecular aspects [71], which enables a more accurate assessment of drug efficacy in a human context; this achievement could contribute to further clinical development of effective disease-modifying therapies.

## 5. Conclusions

PD, which affects over 10 million people worldwide, still remains incurable. The available treatment options are only symptomatic and to date, there is no disease-modifying therapy for PD. Among strategies against α-syn pathology currently under development, the most optimal approach seems to be targeting of toxic oligomers with small-molecules that pass blood-brain barrier, such as anle138b. However, as abrupt disruption of α-syn aggregates could lead to cellular shock and ER stress response, concomitant amelioration of ER stress (e.g. by selective PERK inhibition) could vastly enhance neuroprotection in the course of PD. Here, we evaluated the effect of small-molecule inhibitors of protein aggregation and ER stress in a novel in vitro model of PD, human midbrain organoids generated from iPSC line treated with α-syn PFFs and 6-OHDA. This novel 3D system recapitulates PD pathogenic cascades in a more physiological environment, as compared to 2D cultures or animal models. We demonstrate that combination therapy with anle138b and AMG44 inhibitors significantly increases the PD organoid viability, diminishes the aggregation of pathogenic forms of α-syn, and reduces apoptosis of dopaminergic neurons compared to single therapy with each agent. Mechanistically, the inhibitors fine-tune the expression of α-syn and eIF2α phosphorylation, downregulate pro-apoptotic CHOP factor, and upregulate pro-survival BiP, XBP1s and ATF6 proteins. The results obtained suggest that anle138b and AMG44 or other molecules of similar or even more specific activity could eventually be implemented into clinical trials after extensive preclinical research. We firmly believe that such combined therapy specifically targeting the two molecular events essential to PD pathology may lead to development of ground-breaking treatment strategies against this devastating disorder.

## CRediT authorship contribution statement

Conceptualization, N.S. and I.M.; methodology, N.S.; validation, N.S. and I.M.; formal analysis, N.S. and I.M.; investigation, N.S., M.G., and G.G.; resources, I.M.; data curation, N.S., I.M.; writing—original draft, N.S. and M.G.; writing—review and editing, I.M.; visualization, N.S. and M.G.; supervision, I.M.; project administration, I.M.; funding acquisition, I.M. All authors have read and agreed to the published version of the manuscript.

